# Three-dimensional plant architecture and sunlit-shaded patterns: a stochastic model of light dynamics in canopies

**DOI:** 10.1101/147553

**Authors:** Renata Retkute, Alexandra J. Townsend, Erik H. Murchie, Oliver E. Jensen, Simon P. Preston

## Abstract

**Background and Aims:** Diurnal changes in solar position and intensity combined with the structural complexity of plant architecture result in highly variable and dynamic light patterns within the plant canopy. This affects productivity through the complex ways that photosynthesis responds to changes in light intensity. Current methods to characterise light dynamics, such as ray-tracing, are able to produce data with excellent spatio-temporal resolution but are computationally intensive and the resultant data are complex and high dimensional. This necessitates development of more economical models for summarising the data and for simulating realistic light patterns over the course of a day.

**Methods:** High-resolution reconstructions of field-grown plants are assembled in various configurations to form canopies, and a forward ray-tracing algorithm is applied to the canopies to compute light dynamics at high (1 minute) temporal resolution. From the ray-tracer output, the sunlit or shaded state for each patch on the plants is determined, and these data are used to develop a novel stochastic model for the sunlit-shaded patterns. The model is designed to be straightforward to fit to data using maximum likelihood estimation, and fast to simulate from.

**Key Results:** For a wide range of contrasting 3D canopies, the stochastic model is able to summarise, and replicate in simulations, key features of the light dynamics. When light patterns simulated from the stochastic model are used as input to a model of photoinhibition, the predicted reduction in carbon gain is similar to that from calculations based on the (extremely costly) ray-tracer data.

**Conclusions:** The model provides a way to summarise highly complex data in a small number of parameters, and a cost-effective way to simulate realistic light patterns. Simulations from the model will be particularly useful for feeding into larger-scale photosynthesis models for calculating how light dynamics affects the photosynthetic productivity of canopies.

## INTRODUCTION

Plant canopies are complex three-dimensional (3D) structures in which the light distribution is complicated and dynamic, for example due to solar movement. Diurnal changes in solar position and occlusion caused by overlapping leaves mean that leaves alternate throughout the day between periods in which they are sunlit and periods in which they are shaded. Sun rays that temporarily penetrate to the lower layers of the canopy give rise to ‘sun-flecks’ that are highly intermittent. The spatio-temporal dynamics of direct light influences many fundamental physiological functions, such as photosynthesis, photoacclimation, and photoinhibition (Walters, 2005; Athanasiou et al., 2010; Ruban and Belgio, 2014; Vialet-Chabrand et al., 2017), and secondary biophysical processes such as drought tolerance, water-use efficiency (Qu et al., 2016), plant growth and crop yield (Maddonni et al., 2002). This is because photosynthesis does not directly track the fluctuations in light: for example, delays in photosynthetic induction to high light result from the time taken to activate enzymes in the Calvin cycle, open stomatal pores and build up metabolite pool sizes, and delays in recovery from photoprotection in low light result from the xanthophyll cycle, impacting productivity (Lawson and Blatt, 2014; Retkute et al., 2015; Burgess et al., 2015; Kromdijk et al., 2016; Burgess et al., 2017; Townsend et al. 2018). Optimising photosynthesis by addressing these inefficiencies is clearly a target for improving crop yield but doing so requires a clear understanding of the *dynamic* light conditions in a canopy, rather than just static or time-averaged conditions.

Light dynamics may be measured empirically by various methods, such as hemispherical canopy photographs (Pearcy and Yang, 1996), a photosynthetically active radiation (PAR) sensor moving on a horizontal track (Ross et al., 1998), an electromagnetic 3D digitiser (Sinoquet et al., 1998), or a near-ground imaging spectroscopy system (Zhou et al., 2017). However, spatial resolution of these techniques is typically very poor. This limitation is overcome by digitally reconstructing plants and canopies (Pound et al., 2014) then using ray-tracing to compute light dynamics (Song et al., 2013). In spite of technical challenges, for example due to occlusion and very fine structures such as wheat ears, digital reconstruction of field-grown plants tends to provide a highly accurate description of canopy geometry. However, our understanding of photosynthetic characteristics in canopies is hampered by a current reliance on using ray-tracing to understand the light dynamics in 3D reconstructed canopies (Kim et al., 2016).

Current ray-tracing approaches are costly in computer resources and produce vast data sets as output, especially if computing at high spatio-temporal resolution. Here we develop a novel mathematical model to describe and rapidly simulate sunlit-shaded patterns within a canopy. The model involves two states, sunlit and shaded, and rates of switching between them that we model as functions of time of day and the depth within the canopy. We construct several different realistic digital canopies and use a ray-tracer to identify the times of switching between sunlit and shaded states at positions throughout the canopies. We then use these switching times to estimate the rate functions for switching between states. This offers insight into how light dynamics in a particular canopy depends on time of day and depth within canopy, and how light dynamics varies between canopies involving different plant species, canopy planting density, and canopy leaf area index (LAI).

We use light patterns simulated from the fitted models as an input into a model to predict the reduction in photosynthetic yield attributable to photoinhibition.

## MATERIALS, MODELS AND METHODS

### Digital canopy reconstruction and ray-tracing

To investigate light dynamics in a range of canopies with different structural characteristics, we constructed digital canopies by assembling imaged and digitally reconstructed plants of wheat (lines 1 and 2 in Burgess et al., 2015) and bambara groundnut (from Burgess et al., 2017, at two different growth stages: 39 and 80 days after sowing) in various configurations. The reconstructions represent the surface of a plant with a large number, *N*, of small triangular patches. Figure 1A shows an example of a reconstructed wheat plant, with an individual leaf at the lower part of the plant shown in blue. A triangular patch indexed, say, by *j* is defined by the set of coordinates {**x**_1*j*_,**x**_2*j*_,**x**_3*j*_} of its three vertices. The centroid of this patch is 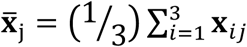, and its normalised height,

**Figure 1.**
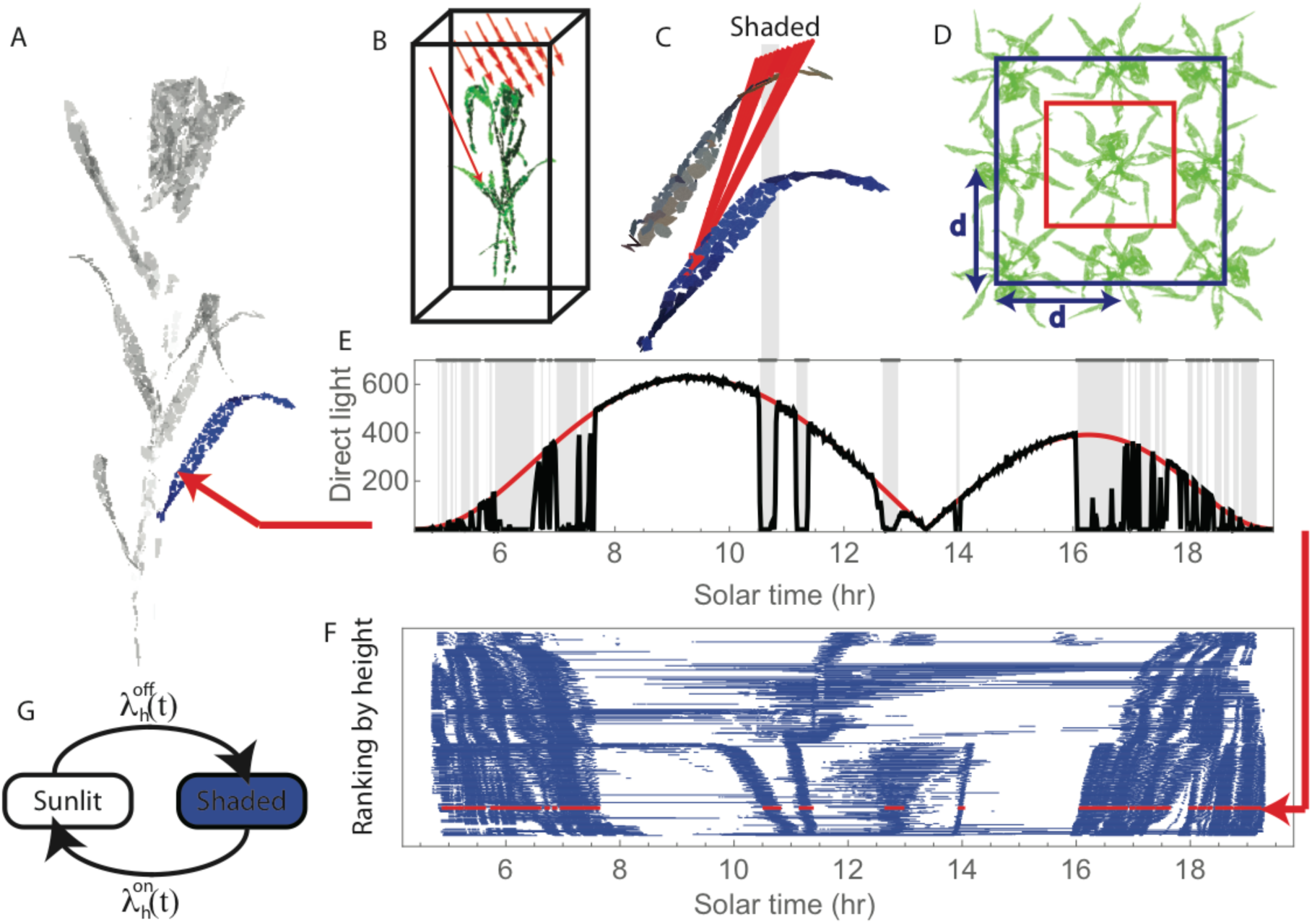
Quantifying sunlit and shaded dynamics. (A) Reconstructed wheat plant from Burgess et al., 2015. (B) Set-up for the ray-tracer (Song et al., 2013): red arrows show direct light rays; once a ray hits the boundary of the bounding box, it is moved to the opposed vertical face of the box. (C) Shading will occur when a ray is obstructed by other leaves or a stem. (D) Construction of canopies was done in two ways: putting the bounding box just outside the plant (red rectangle) or putting plants on a 3 × 3 grid at a distance d apart and putting the bounding box through the centres of boundary plants (blue rectangle). (E) Diurnal dynamics of light, in µmol m^2^s-^1^, at a particular patch showing ray-tracer simulation (solid black curve), light amplitude envelope (solid red curve) and inferred shaded periods (horizontal grey lines). Time resolution is 1 minute. One of the shaded periods is extended to (C) to indicate schematically the occlusion caused. (F) Sunlit-shaded patterns for the patches comprising the leaf shown in blue in (A); each row corresponds to an individual patch, with patches ordered by the height of their centroids. The row shown in red corresponds to the particular patch shown in (E). (G) The two-state sunlit-shaded model: switching on (from shaded to sunlit) occurs at rate 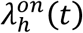 and switching off (from sunlit to shaded) at rate 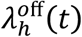.

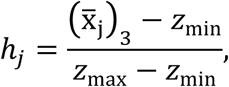

in which 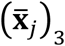 denotes height from the ground of the *j*th patch’s centroid, and z_min_ = min_*ij*_{(**x**_*ij*_)_3_}, z_max_ = max_*ij*_{(**x**_*ij*_)_3_} are respectively the minimum and maximum heights amongst all the vertices in the canopy. The models developed later involve dependence on these normalised heights.

We constructed canopies *in silico* by arranging into various configurations several individual-plant reconstructions of Burgess et al., 2015, and Burgess et al., 2017, exploiting the periodic boundary conditions of the ray-tracer (explained below) which give a natural way to “tile” individual plants to form an effective canopy. We investigated two ways to do this: (i) by putting the bounding box just outside the plant (as shown by the red rectangle in Figure 1D); or (ii) arranging plants on 3 × 3 square lattice a distance *d* apart, and putting the bounding box through the centres of boundary plants (as shown by the blue rectangle in Figure 1D). The periodic boundary conditions mean that case (i) amounts to considering an infinite square lattice of identical copies of the same plant. Case (ii) is similar, but it introduces additional heterogeneity through randomising the orientation of the different plants. We positioned the plants at distances *d* equal to 200mm, 150mm, 125mm and 100mm. In case (ii), we analysed the light dynamics for the plant in the centre of the 3 × 3 lattice. Configuring plants in these various ways led to canopies with a wide range of different structures and LAI.

To compute the light distribution within the constructed canopies we used *Fast-Tracer v3.0* (Song et al., 2013), a software implementation of a forward ray-tracing algorithm. This simulates three categories of light (direct, diffuse and scattered light), and determines where individual rays of light are eventually absorbed on leaf surfaces. Figure 1B shows a configuration for the ray-tracer software (Song et al., 2013). In-bound rays are arranged over a grid above the plant. The direction and amplitude of each ray depends on latitude and time of day. Ray tracing is performed in a cubic domain with periodic boundary conditions on the vertical faces so that when a ray exits one boundary of the domain it re-enters on the opposite vertical face. We used latitude 53° (for Sutton Bonington, UK), atmospheric transmittance 0.5, light scattering 7.5%, light transmittance 7.5%, day 182 (1 July), corresponding to the location that the plants were grown and the day they were imaged (Burgess et al., 2015). We calculated the direct light intercepted during the day at 1 minute resolution for every patch in the canopy. The high temporal resolution enabled us to investigate even short-term light fluctuations in the canopy. Figure 1E shows an example of the light pattern computed for a particular patch.

### Computing sunlit-shaded patterns from ray-tracing data

To construct sunlit-shaded patterns for each patch we compared values of direct light computed by the ray-tracing algorithm (Song et al., 2013) to the direct light irradiance, *A*dr, on a unit surface in the absence of any shading (Cambell and Normal, 1998); the latter, defined in Equation (18) in the Appendix, depends on latitude, day of year, time of day, and the angle between a light ray and the normal to the patch in question. Figure 1E shows, for a particular patch in the lower part of a canopy, the direct light computed from the ray-tracer (in black) and *A*dr (in red). We designated a patch at a given time point as being shaded if the value of direct light computed by the ray-tracer differed from *A*dr by more than 10%. The shaded periods are indicated in Figure 1E by vertical grey bars. The substantial shaded period between 10 and 11 o’clock, for example, shown in Figure 1E is a consequence of the shading shown in Figure 1C. These binary sunlit-shaded light patterns, computed for each patch in the canopy, are the inputs to the models we develop below. Figure 1F shows the sunlit-shaded patterns for all the patches comprising the leaf shown in blue in Figure 1A, with each row corresponding to an individual patch, the patches (and hence rows) having been ordered according to the normalised heights of the patch centroids. The diagram reveals an intricate pattern, with shadows from the upper leaves moving along the surface of the leaf as the sun changes position in the sky.

In the following sections we develop models for the sunlit-shaded patterns: first Model 1, a simple preliminary model which we use to introduce ideas and notation; and then Model 2, which is the novel modelling contribution of this paper. In each case, we present (i) the model definition, (ii) how the model can be fitted to experimental data, and (iii) how the fitted model can be simulated to generate realistic light patterns.

### Sunlit-shaded dynamics: Model 1

In this initial model, we limit attention to a single patch and consider how its rate of switching from sunlit to shaded, or vice versa, changes with time, *t*, in a time interval of interest, (0,*T*). The central assumption is that switching events arise from a non-homogeneous Poisson process (see e.g. Ross, 2006). A non-homogeneous Poisson process is a stochastic process defined via: (i) an intensity function *λ(t) ≥ 0* such that, for *0 < δt ≪ 1*,

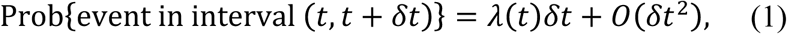

where *O(δt^2^)* denotes terms involving squared or higher powers of *δt* which are negligible for small *δt*; and (ii) the assumption that the probabilities of events in distinct intervals are independent (Ross, 2006). From this independence, and Equation (1), it follows that for any interval *(t, t + u)⊆* (0,*T*),

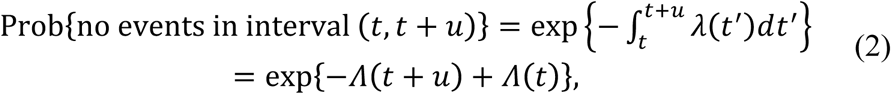

where 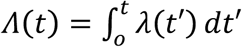 Equation (2) is useful in the following section for constructing expressions needed for fitting the model to data.

### Fitting Model 1

The goal is to fit the model by estimating the intensity function *λ(t)* based on a set of switching times *0 < v_1_ <…<v_n_<T*. We will use maximum likelihood estimation, a standard statistical principle for estimating model parameters from data (Cox, 2006). This involves constructing the likelihood function for the model, which is the probability (density) function evaluated at the observed realisation of switching times but regarded as a function of the parameter *λ(t)* to be estimated. The likelihood function for this model is

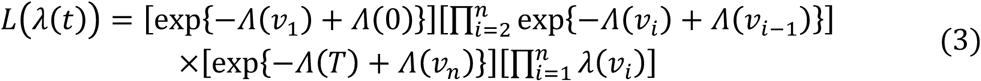

can be derived by discretising the interval (0, *T*) with increments of size *δt*, writing the likelihood as a product of factors (using independence of increments) with each factor either (1) or its complement, depending on whether the increment contains an event, then taking the limit *δt → 0*. The four factors in square brackets have the following interpretations: the first factor is a contribution from having no events in the interval *(0, v_1_),* the second factor from having no events in *(v_i_, v_i_-1)* for *i = 1,…, n*, the third from having no event in the interval *(v_n_, T)*, and the fourth is the contribution from the switching events occurring at times *v_1_,…, v_n_*. These interpretations are helpful in constructing likelihood functions for Model 2 below, but in the present case, telescoping in the exponent means that (3) simplifies to

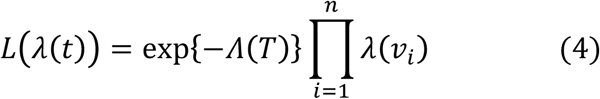

Maximising *L(λ(t))* directly with respect to an unrestricted *λ(t)* is ill-posed (since the maximising *λ(t)* would blow up at the switching instants *t =v_1_,…, v_n_*, and be zero elsewhere). A solution to this is to impose a functional form for *λ(t)* in terms of a small number of parameters, ***θ*** *= (θ_1_,…, θ_p_).* We then write *λ(t) = λ(t;* ***θ***), and fit the model by maximising the likelihood (4) with respect to ***θ***. In fact, it is equivalent and usually more convenient to compute this maximum likelihood estimate (MLE) of ***θ*** by maximising the log of the likelihood function, which is

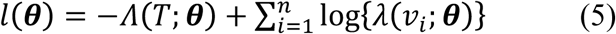

We discuss below specific choices for the form of *λ(t;****θ***). Function (5) can be maximised by a numerical optimisation routine, nd for the calculations in this paper we have used the Nelder–Mead simplex method (Nelder and Mead, 1965). If *λ(t;****θ***) is linear in ***θ*** then (5) is concave in ***θ*** making the numerical optimisation particularly straightforward.

### Simulating from Model 1

From (1), the distribution function for the additional time until the next event occurs given that an even occurred at time *v* is

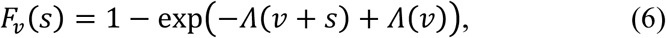

and a random variable can be simulated from this distribution using the inversion method (Ross, 2006). An algorithm to simulate a sequence of event times *v_1_, v_2_, v_3_,*… is thus as follows. Let *v_1_* be a simulated value from the distribution *F_0_*. Then let *v_2_* equal *v_1_* plus a simulated value from the distribution F_*v1*_. Continue in this way, letting *v_i+1_* equal *v_i_* plus a simulated value from the distribution F_*vi*_ until *v_i+1_ > T*.

### Sunlit–shaded dynamics: Model 2

The main contribution of this paper is to extend Model 1 in two ways: (i) to incorporate distinct rate functions, *λ*^*on*^(*t*) and *λ*^*off*^(*t*), for switching “on” (from shaded to sunlit) and “off” (from sunlit to shaded), respectively; and (ii) to describe multiple patches, with the rate functions for different patches depending on the normalised height, *h*, within the canopy (in addition to time, *t*, as in Model 1).

Extension (i) requires a notational distinction between the times of on-switching events, say *xi*, and off-switching events, *y_i_*. For a given patch, on-and off-switching events necessarily alternate, and hence a sunlit-shaded pattern is characterised by the ordered set of times *{x^1^, y_1_, x_2_, y_2_,…, x_n_, y_n_}.* We represent a state initially “on” at time 0 by having *x*_1_<0, and “off” at time *T* by *y*_*n*_ > *T* (the particular values of *x*_1_ and *y*_*n*_ in these cases do not need to be specified) but besides these exceptions we otherwise assum that *0<x_i_<y_i_<_T_* for all *i*. Figure 2 illustrates the notation, with the four different examples showing the four possible cases involving the different combinations of “on” and “off” states at *t = 0* and *t = T*.

**Figure 2.**
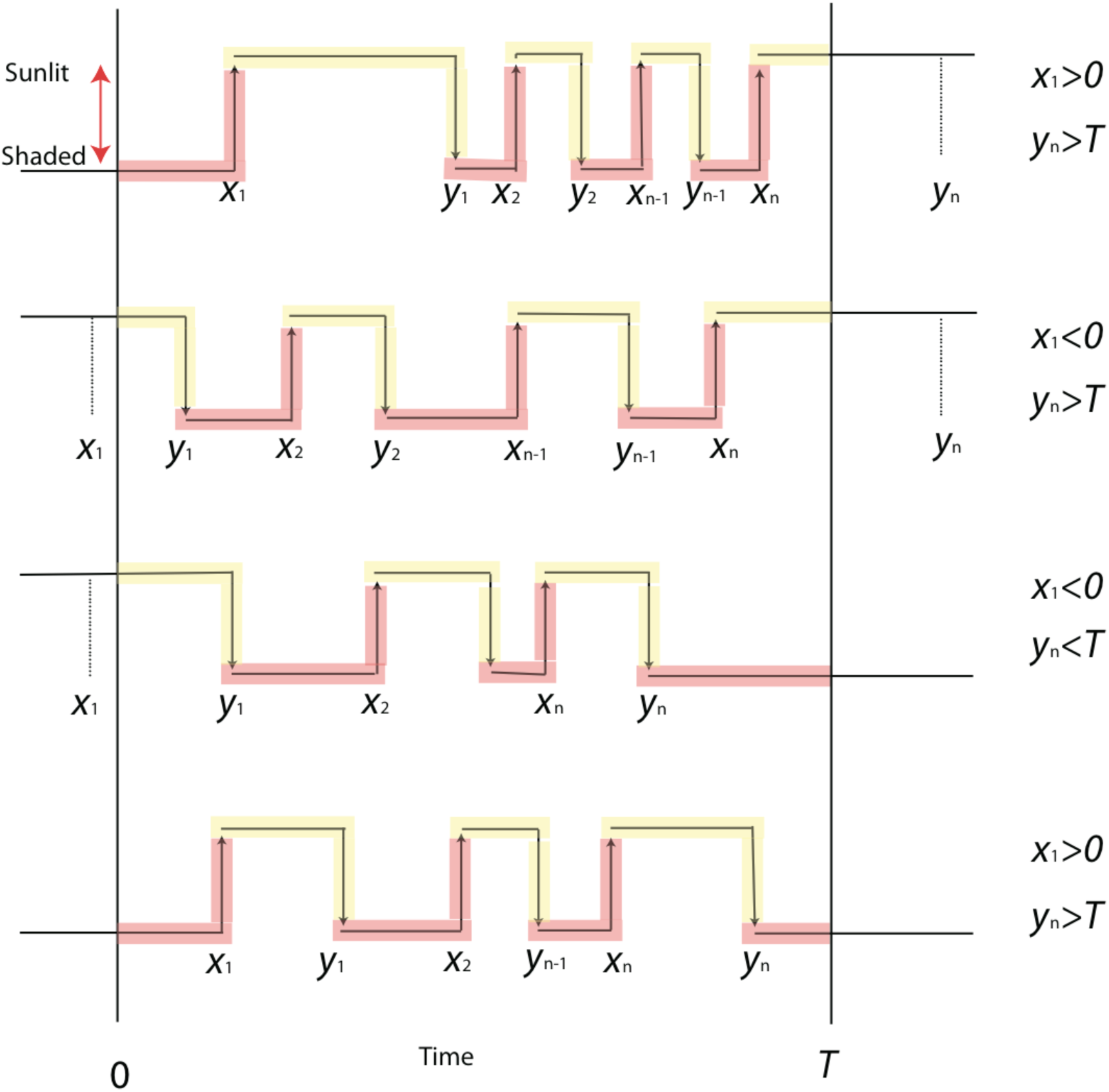
Illustration of the notation for the model, showing the four possible combinations of states at the beginning and end of the interval [0, *T*]. At time *t*=0, a sunlit state is indicated by *x*1 < 0 and a shaded state by *x*1 > 0; at time *t*= *T* a sunlit state is indicated by *yn* > *T* and a shaded state by *yn* > *T* The different sections are coloured to indicate how they contribute to the (log) likelihood functions: red denotes a contribution to on-switching functions (7, 9) and yellow to off-switching function (8, 10).

### Fitting Model 2

In terms of the switching times, *{x_1_, y_1_, x_2_, y_2_,…, x_n_, y_n_},* for a given patch, the likelihood functions for *λ*^*on*^(*t*) and *λ*^*off*^(*t*) are then

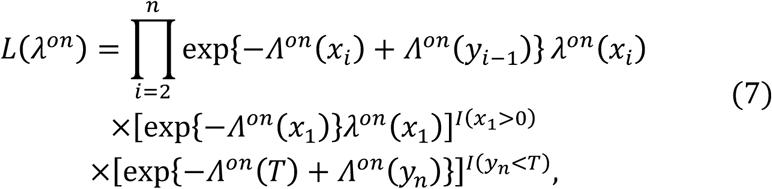

and

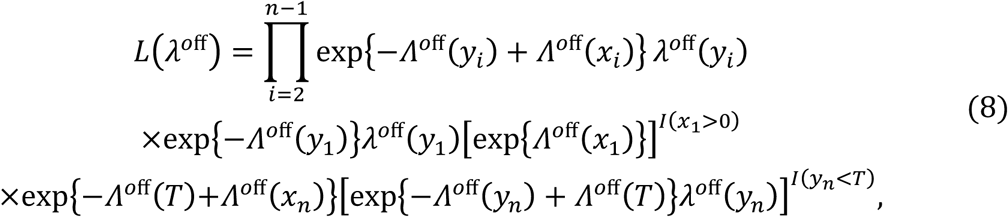

where *I(·)* is the indicator function, equal to one if its argument is true and zero otherwise. Equations (7) and (8) generalise (3) to distinguish between sunlit-to-shaded and shaded-to-sunlit switches, and they are constructed in a similar way to (3); see Figure 2 in which sections of example sunlit-shaded patterns are coloured to indicate how they contribute to either (7) or (8).

The final step is to generalise to multiple patches, incorporating dependence of the rates on the heights of the different patches. Let *j = 1,…, m* index the different patches, and quantities specific to the *j*th patch be indicated by suffix *j*. As before, we assume *x*_1,*j*_*<0* and *y*_*n,j*_ > *T* if the state is “on” at the beginning and end, respectively, of the interval (0, *T*). We let *h*_*j*_ denote the height of the *j*th patch and use subscripts on the rate functions to denote their dependence on height, i.e., the rate functions for the *j*th patch are *λ*^*on*^*(t)* and *λ*^*off*^*(t)*. Assuming independence of patches (an assumption discussed later in the Discussion section), the likelihood functions for *ffon(t)* and *ffoλ(t)* can be constructed as a product of factors of the form (7) or (8) over index *j = 1,…, m*, giving log-likelihood functions

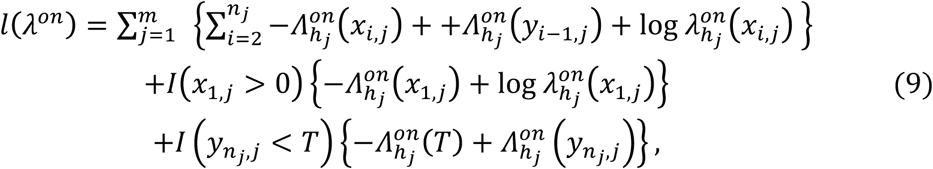

and

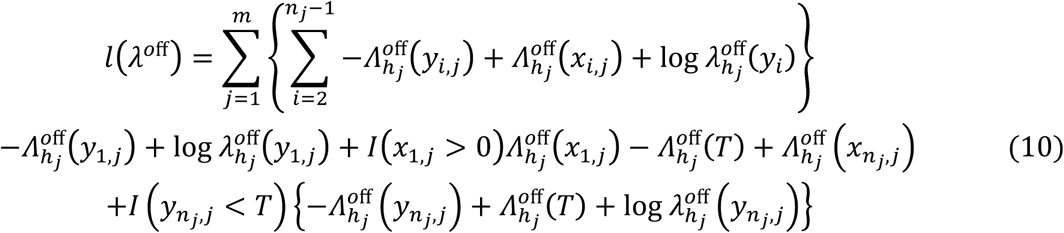

Like before, it is necessary to choose functional forms for *λ*^*on*^_*h*_*(t)* and *λ*^*off*^_*h*_*(t)*, and we discuss specific choices in the Results sections below.

### Simulating from Model 2

This model distinguishes whether at time *t = 0* a patch is in a sunlit or shaded state.

For simulations we choose a random starting state from the distribution

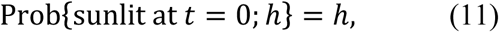

where *h* is the normalised height of the patch’s centroid, so that patches high in the canopy tend to start off sunlit whereas those at the bottom tend to start shaded.

The distribution for the time until the next event depends on whether switching is from sunlit to shaded or vice versa. We denote the distribution for the time to the next “on” event given an “off” event occurred at time *x*1 by 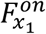; and the time to the next “off” event, given an “on” event occurred at time *y*_1_ by 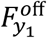. An algorithm to simulate from Model 2 is then as follows. Simulate the initial state as either sunlit or shaded. Supposing it is sunlit, let *x*_1_ be a simulated value from 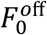 Then let y1 equal *x1* plus a simulated value from 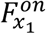. Continue letting *x*_*i+1*_ equal *y*_*i*_ plus a simulated value from 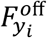 and *y*_*i+1*_ equal *x*_*i+1*_ plus a simulated value from 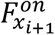 until either *x*_*i*+1_ > *T* or *y*_*i*+1_ > *T* If in the first step above the initial state is instead shaded, then the following two steps are replaced by simulating *y*_1_ from 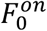, but the algorithm otherwise proceeds the same.

This algorithm simulates the binary state of whether the patch is sunlit at time *t*. The corresponding direct light flux density (adjusting intensity during sunlit periods to account for factors including solar position at time *t* and patch orientation) is

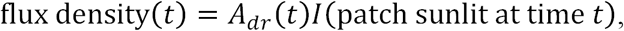

where Adr is as described earlier and defined in (18) in the Appendix.

### Case study: Photoinhibition model

To assess the models, we investigate whether light patterns simulated from Model 2 and used as input into a photoinhibition model lead to similar results compared with when the ray-tracer dynamics are used as input. Photoinhibition is a light-dependent decline in the maximal quantum yield of photosynthesis and can lead to a lowering of photosynthesis and potential growth (Long et al., 1994) and hence it is a good physiological quantity with which to test the impact of light dynamics. The effect of photoinhibition can be characterised by changes in the shape of the light-response curve, in terms of changes in the parameters that define it. The light-response curve is often modelled by a non-rectangular hyperbola (defined in the Appendix) involving two shape parameters: the quantum yield of PSII, *φ*, and convexity, *θ*.

Following Burgess et al., 2015, we quantify the impact of photoinhibition by predicting the reduction in carbon gain over a day within a wheat canopy. We use the same canopy, photoinhibition model and physiological measurements as Burgess et al., 2015, so that the only difference here is that the light dynamics are simulated from the model, rather than coming directly from the ray-tracer. The model of photoinhibition was parameterised by field data consisting of chlorophyll fluorescence and light-response curves of carbon dioxide assimilation. The canopy was divided into three layers (top, middle and bottom), and for leaves at each layer light-response curves and dark-adapted maximum quantum yield, *F*_*v*_/*F*_*m*_, were measured at a midday, giving scaling factors 0.857 for the top layer and 0.955 for the middle layer.

## RESULTS

### Model 1: example of simulation and model fitting

For *λ*(*t*) we assume the simple functional form *λ*(*t*) = 3 + 0.05*t* – 0.075(*t* – 6)^2^. Here time *t* is measured in hours over a *T=12*hr period starting at 6am, so that *t=t*_*md*_*=6*hr represents noon. The coefficients have been chosen somewhat arbitrarily to simulate a high degree of variation over the period, in which the rate of switching increases from sunrise, reaches its maximum near the middle of the day, then decreases until sunset. Figure 3B shows plotted in grey, and Figure 3A shows a single realisation from this model.

**Figure 3.**
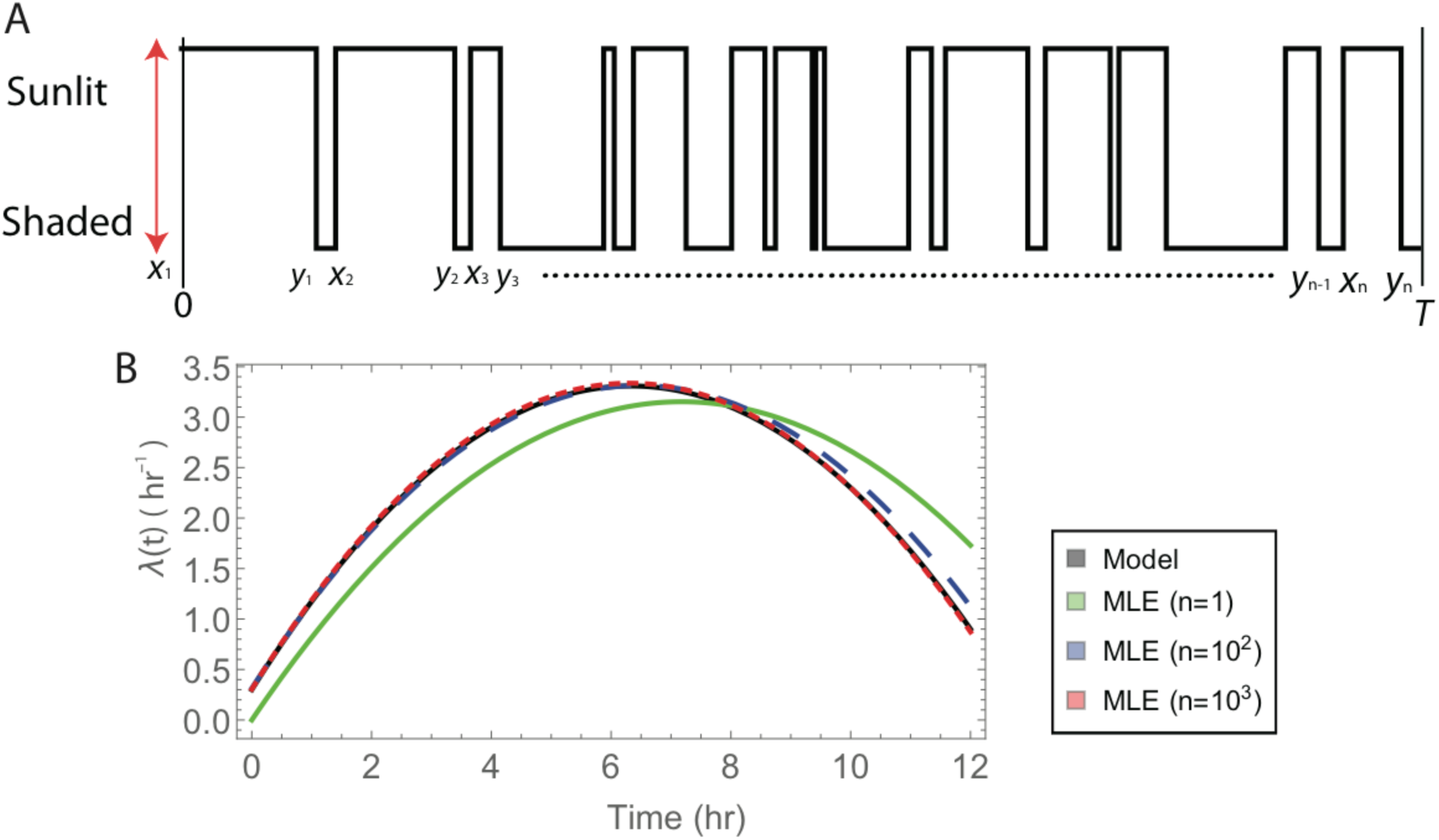
A simulated realisation of Model 1, and maximum likelihood estimation from fitting the model to the realisation. (A) A realisation with *λ*(*t*) = 3 + 0.05*t* – 0.075(*t –* 6)^2^. (B) A plot of this true *λ*(*t*) together with estimates of it based on from data with various numbers of realisations, *n*, showing convergence of the estimates to the true *λ*(*t*) as *n* grows.

Using maximum likelihood estimation to fit the model *λ*(*t*; *θ*) = *θ*_1_ + *θ*_2_*t* + *θ*_3_(*t −* 6)^2^ based on the realisation in Figure 3A leads to the estimated intensity function 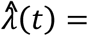 (shown in green in Figure 3A). The estimated intensity function matches reasonably well with the true intensity function, but not exactly because the estimate is based on a small amount of data. Typically, the MLE gets closer to the true answer as the amount of data increases. This is illustrated by the blue and red curves in Figure 3B, which are the MLEs based on data from multiple realisations.

### Model 2 fitted to ray-tracer data

We constructed canopies by assembling 3D reconstructions of wheat and Bambara groundnut plants, as described above. Figure 4 shows images of plants and canopies we constructed and analysed: an individual wheat plant in 4 different orientations; the plant randomly rotated and positioned at distances *d* = 200mm, 150mm, 125mm, and 100mm from a replica of a plant from the same line; individual plants from different wheat lines; and individual Bambara groundnut plants at two different growth stages (39 and 80 days after sowing). Canopies (A)-(D) and (I)-(J) are configurations for which we placed the bounding box just outside the plant (as shown by red rectangle in Figure 1D), whereas for canopies (E)-(H) we arranged plants on a 3 *×* 3 square lattice at different distances *d* apart. Cumulative leaf area index (cLAI), shown in Figure 4(ii) for the various canopies, describes how plant mass accumulates from top to bottom of each canopy. The cLAI profiles for canopies (A)-(D) are identical since rotation of the plant does not change the distribution of leaf mass with respect to depth.

**Figure 4.**
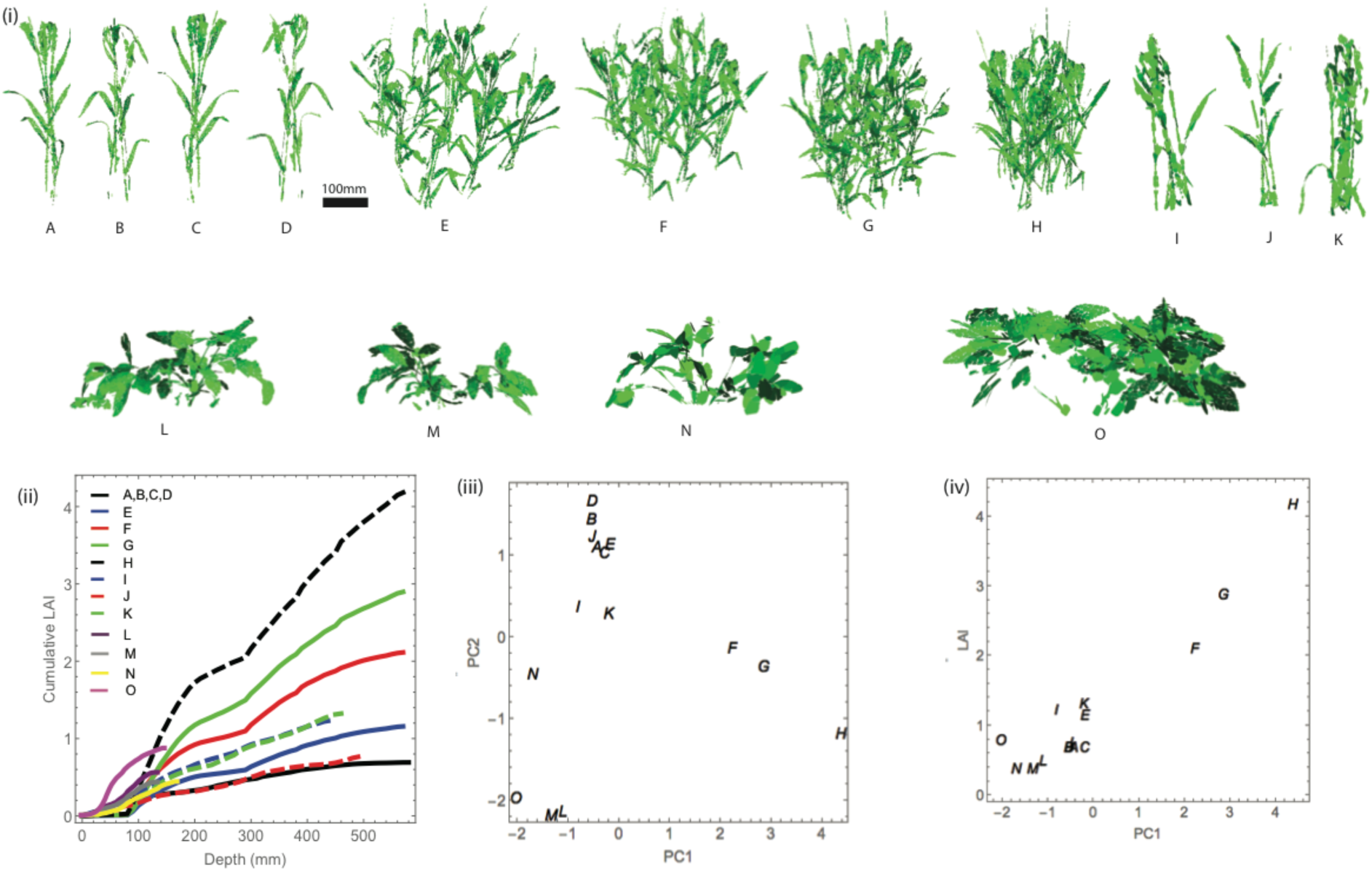
Plants, canopies, and fitted models: (i) Reconstructed plants (**A-O**), (ii) cumulative leaf area index as a function of depth, (iii) principal component analysis of fitted parameters, and (iv) relationship between the first principal component and LAI. Images of original plant (**A**); original plant rotated 90° (**B**), 180° (**C**), and 270° (**D**); original plant randomly rotated and positioned at distances 200mm (**E**), 150mm (**F**), 125mm (**G**), and 100mm (**H**); replica of a plant from the same line (**I**); plants from two different wheat lines (**J** and **K**); three plants of Bambara groundnut 39 days after sowing (**L, M, N**) and 80 days after sowing (**O**). For a detailed description of the lines and reconstructions, see Burgess et al., 2015, and Burgess et al., 2017.

For the rate functions 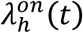 and 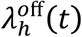 we chose

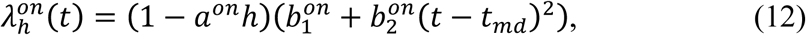

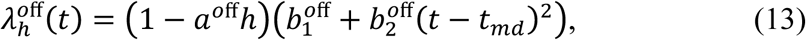

so that the switching rates depend linearly on height, *h*, and parabolically on *t*. This is the simplest form we can choose for (12) and (13) such that they depend on *h* and *t*, and with their dependence on *t* being both smooth and symmetric about midday.

We estimate the parameters 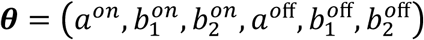 from the sunlit-shaded patterns extracted from ray-tracing data by maximum likelihood estimation, as described above. Table 1 shows the fitted parameters for the various canopies we considered, and each canopy’s LAI. Notable from the table is that *a*^*on*^ and *a*^*off*^ both tend to be far from zero, indicating that the switching rates are strongly dependent on depth within the canopy. For many of the canopies *a*^*off*^ is very close to 1, so that the off rate at the very top of the canopy is close to zero; this is because patches at the very top are not obstructed by other leaves and are hence permanently sunlit. Typically, the parameters 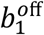 and 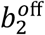 contributing to the off rate are larger for canopies with higher LAI and the corresponding parameters 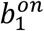 and 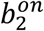 in the on rate are slightly smaller. This is consistent with the intuition that in dense canopies sun flecks are typically shorter and less frequent.

**Table 1.**
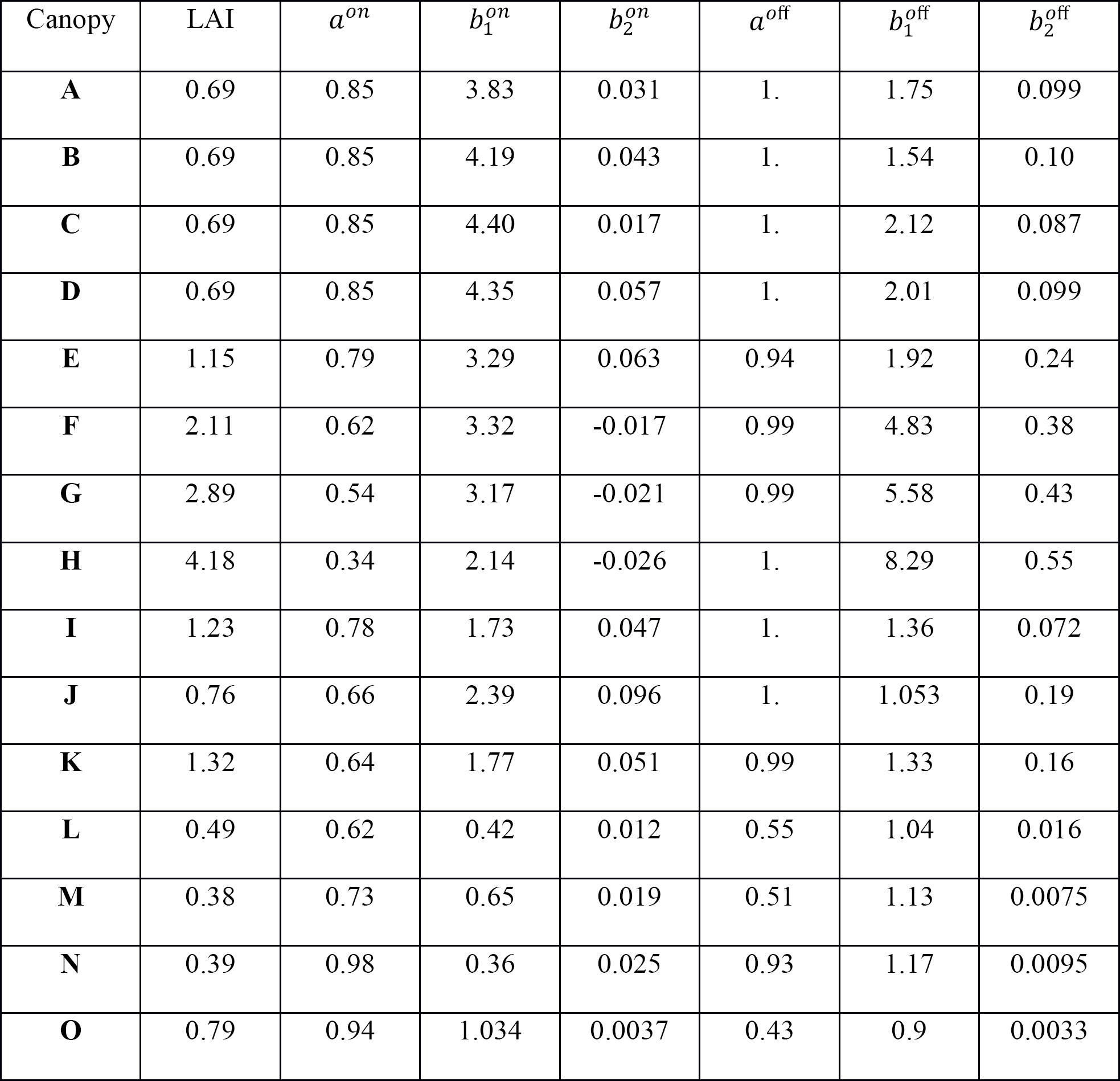
Parameter estimation for the model (Eq.(12)-((13). The canopy letters correspond to canopy labels shown in Figure 4.

| Canopy | LAI | $a^{on}$ | $b_1^{on}$ | $b_2^{on}$ | $a^{off}$ | $b_1^{off}$ | $b_2^{off}$ |
| --- | --- | --- | --- | --- | --- | --- | --- |
| **A** | 0.69 | 0.85 | 3.83 | 0.031 | 1. | 1.75 | 0.099 |
| **B** | 0.69 | 0.85 | 4.19 | 0.043 | 1. | 1.54 | 0.10 |
| **C** | 0.69 | 0.85 | 4.40 | 0.017 | 1. | 2.12 | 0.087 |
| **D** | 0.69 | 0.85 | 4.35 | 0.057 | 1. | 2.01 | 0.099 |
| **E** | 1.15 | 0.79 | 3.29 | 0.063 | 0.94 | 1.92 | 0.24 |
| **F** | 2.11 | 0.62 | 3.32 | -0.017 | 0.99 | 4.83 | 0.38 |
| **G** | 2.89 | 0.54 | 3.17 | -0.021 | 0.99 | 5.58 | 0.43 |
| **H** | 4.18 | 0.34 | 2.14 | -0.026 | 1. | 8.29 | 0.55 |
| **I** | 1.23 | 0.78 | 1.73 | 0.047 | 1. | 1.36 | 0.072 |
| **J** | 0.76 | 0.66 | 2.39 | 0.096 | 1. | 1.053 | 0.19 |
| **K** | 1.32 | 0.64 | 1.77 | 0.051 | 0.99 | 1.33 | 0.16 |
| **L** | 0.49 | 0.62 | 0.42 | 0.012 | 0.55 | 1.04 | 0.016 |
| **M** | 0.38 | 0.73 | 0.65 | 0.019 | 0.51 | 1.13 | 0.0075 |
| **N** | 0.39 | 0.98 | 0.36 | 0.025 | 0.93 | 1.17 | 0.0095 |
| **O** | 0.79 | 0.94 | 1.034 | 0.0037 | 0.43 | 0.9 | 0.0033 |

The information in Table 1 can be visualised in two dimensions using Principal Component Analysis (PCA). We performed PCA based on the correlation matrix of the fitted 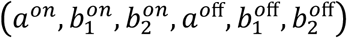 parameters. The first PC, which explains 59% of the variability, has loadings (0.27, 0.54, 0.54, −0.46, 0.18, −0.30) suggesting the interpretation that it contrasts the off and on rates, or in other words that it is a measure of typical “shadedness” within the canopy. Indeed, the first PC correlates very strongly with LAI (Pearson correlation coefficient of 0.984, p-value<10-^7^); see Figure 4(iii). The second PC, which explains 30% of the variability, has loadings (0.60, −0.09, 0.04, 0.2, 0.60, 0.49), with the dominant values corresponding to *a*^*on*^,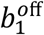 and 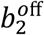. This has the interpretation of contrasting the height-dependent part of the on rate with the constant and time-dependent parts of the off rate. Taking this together with Figure 4(iii), which shows clear clustering in PC2 according to species, lines and planting density, this PC provides insight into how the light dynamics between different canopies differ by depth and time of day. Plotting PC1 versus PC2, as in Figure 4(iii), indicates very clear clustering of canopies we expect to be similar, showing clearly that the PCs (and the fitted parameters from which they were computed) encode meaningful information about the canopies.

### Model evaluation: summary plots and photoinhibition case study

To assess the goodness of fit of Model 2, we show in Figure 5 a comparison of the distributions of sunlit and shaded periods, aggregated by canopy over *h* and *t*. The periods for the ray-tracer data tend to be slightly more variable than for the fitted model, which is consistent with our imposing, via (12) and (13), stringent smoothness in the dependence of rates on h and t (and imposing that there is no dependence at all on the other spatial coordinates) which restricts variability between patches. This aside, the histograms match well.

**Figure 5.**
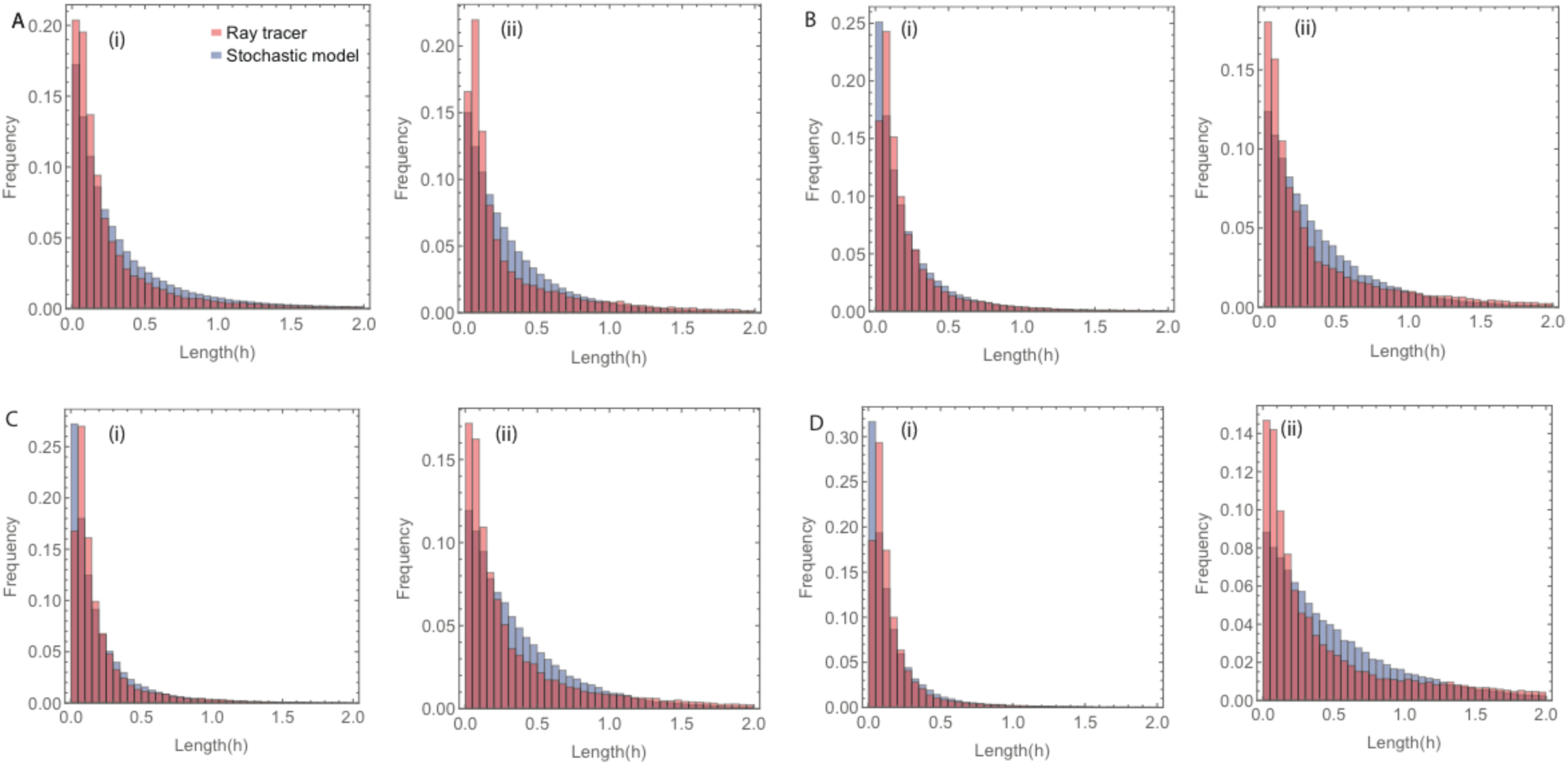
Distributions of the duration of sunlit and shaded periods. Results shown are from ray-tracer (red) and stochastic model (blue) simulations for canopy **E** with plants at distance 200mm (A), canopy **F** with plants at distance 150mm (B), canopy **G** with plants at distance 125mm (**C**) and canopy **H** with plants at distance 100mm (**D**). Panels show distributions of length for (i) sunlit periods, and (ii) for shaded periods. Parameters are as given in Table 1.

As a further evaluation, we consider how different the outcome is if we feed into a photoinhibition model light dynamics simulated from Model 2, rather than from the ray-tracer. To do this, it is necessary to estimate diffused light values, as the rate of photosynthesis depends on the total intercepted irradiance. In contrast to direct light and its properties discussed above, the diffused light does not fluctuate during the day, but depends on the position of a patch within a canopy, latitude, day of a year, and time of the day. The latter three attributes (latitude, day and time) can be used to calculate the diffused light profile over a day on a horizontal surface. Diffused light over all patches has the same shape as this profile, but each patch has an individual scaling of diffused light amplitude. Figure 6A shows profiles of diffused light during a day on a particular patch obtained from the ray-tracer (red) and by fitting a scaling factor to the analytic expression of direct light (given in the Appendix). We have fitted scaling factors using a least squares method to all patches of Line 2 in Burgess et al., 2015. This is shown in Figure 6B as grey dots. To determine the scaling-factor dependence on the normalised height, we calculated an average value of the scaling factor in intervals [*i*/100,(*i* + 1)/100], *i* = 1,…,99; (black curve in Figure 6B).

**Figure 6.**
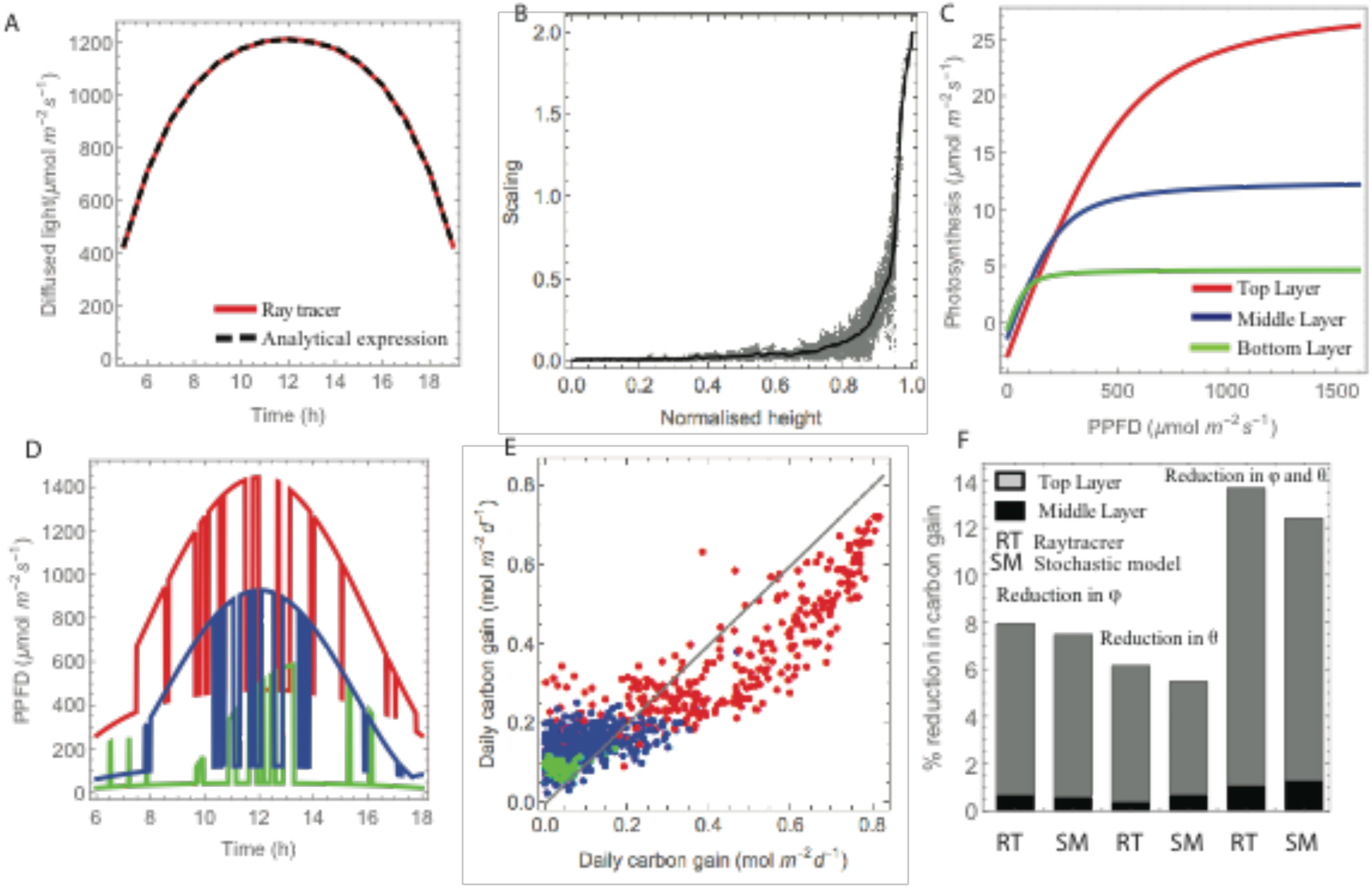
Using model simulations to study photoinhibition. (A) A diffused light profile for a patch at the top of a canopy. (B) Scaling of diffused light as a function of normalised height (gray dots) and fitted spline. (C) Light-response curves for the top (red), middle (blue), and bottom (green) layers (Burgess et al., 2017). (D) Simulated light patterns, and (E) calculated daily carbon gain based on ray-tracer (x-axis) and emulator (y-axis) (with colours matching (C)). (F) The predicted effect of photoinhibition on carbon gain based on ray-tracer (RT) Burgess et al., 2016, and stochastic model (SM)

Photosynthetic rate is light-intensity dependent and so depends on the position of the patch, and the light response curves were measured at the top, middle and bottom of the canopy (Burgess et al., 2015). The maximum photosynthetic capacity was estimated as 28.6µmol m-^2^s-^1^ for the top layer, 12.6µmol m-^2^s-^1^ for the middle layer and 4.7 µmol m-^2^s-^1^ for the bottom layer. Light response curves were taken from Burgess et al., 2015; see Figure 6C. As the LAI of the canopy from Line 2 in Burgess et al., 2015; was close to the LAI of the canopy G, we used parameter values from Table 1 associated with this canopy. We simulated light patterns for all patches for the Line 2 canopy (Burgess et al., 2015) with direct light calculated using simulations from Model 2 and diffused light calculated using the relationship of the scaling factor and normalised height as discussed above. A few of the light patterns from different layers are shown in Figure 6D.

First, we compared daily carbon gain calculated based on light patterns from both model simulations and ray-tracer output for 1000 patches selected uniformly at random; see Figure 6E. The values plotted on the vertical axis are averages over 10 realisations of sunlit-shaded patterns simulated from the model. The results show strong correlation for the two sources of light patterns (Pearson correlation coefficient 0.92, p-value< 10-^6^). Departures from linearity in this plot result from our assuming, via (12) and (13), a very smooth dependence of the switching rates on height, h, and making no attempt to characterise heterogeneity amongst patches at common *h*; the model hence characterises an “average” patch at each height *h* such that, for example, there are no predictions of patches with exactly zero carbon gain (unlike for the ray-tracing patterns in which some patches remain permanently shaded). As with any model, the model could be made arbitrarily more complex to explain more of the variability in the data (perhaps with addition of random effects; see e.g. Dunson, 2008), but here we favour keeping the model simple and interpretable, and in view of the model involving only 6 parameters, the agreement shown in Fig 6E is very good. See the Discussion section for more on the topic of model complexity.

Finally, we have used light patterns obtained using the stochastic model to infer the effect of photoinhibition. We analysed three scenarios: reduction in quantum use efficiency, *φ*, reduction in the convexity, *θ*, and reduction in both *φ* and *θ*. It has been shown previously in Burgess et al., 2015, that the latter scenario gives the largest reduction in carbon gain relative to a noninhibited canopy, and this reduction mostly comes from the top layer. Results in Figure 6F show generally good agreement between using the simpler stochastic model and using the full ray-tracing data, as in Burgess et al., 2015, in predicting the reduction in carbon gain. In the top layer the reduction is consistently slightly lower than for the ray-tracer, but this is actually more in line than Burgess et al., 2015, with other photoinhibition studies, e.g. Zhu et al., 2004.

## DISCUSSION

We have used ray-tracing to compute the light dynamics in complex canopies and developed a novel model to characterise the dynamics. The model is useful for summarising vast and complex ray-tracing data in a small number of parameters, and for simulating light dynamics in a simple and computationally cheap way. Comparing fitted models offers a way to understand differences in light dynamics between different canopies, and the models can be easily simulated to generate realistic light patterns to use as inputs to larger-scale models, for example for computing absorbed radiation and photosynthetic production of a canopy.

In the new field of plant and crop phenotyping, high resolution 3D canopy reconstructions can now be developed routinely, but bottlenecks exist in analysing them for physiological function. For example, using the reconstructions available in Pound et al., 2014, running the ray-tracer “Fast-Tracer” to provide data from a wheat canopy (9 plants) for a simulated 24-hour period can take several days. In comparison the stochastic model takes less than a minute to simulate an individual direct light pattern without the need to run calculations for all of the canopy. Light dynamics characterised by the model are a means to investigate canopy photosynthetic responses (as in Figure 6) and various aspects of crop cultivation such as varietal selection and altered architectural characteristics, and cultivation practice such as cropping system, row spacing, etc.

In this paper we have fitted the models based on ray-tracing data, and so have not avoided the computational cost of ray-tracing. However, in work not presented here we have also investigated fitting the model to only a small random subset of the patches and have found that models fitted this way typically do not differ much from the full fitted models. For the small subset of patches, sunlit-shaded patterns can be computed by simple geometrical reasoning (considering whether there is line-of-sight between a patch and the sun as the day progresses), sidestepping ray-tracing altogether. This is a highly promising for making the model fitting very fast, and thus opening possibilities for using the model for high-throughput analysis.

As with any model, our model is only an abstraction, intended to be a simple description of something complex, which retains only the features of greatest importance at the expense of discarding others. We make no attempt, for example, to describe spatial correlation between the light patterns of different patches. There seems no obvious way to do so without retaining the full 3D geometry of the canopy, and this would forsake the simplicity that makes the model useful. In any case, we do not foresee many applications of a light-dynamics model requiring such high spatial resolution that spatial correlation is important. The assumption of independent patches, made in constructing (9) and (10), embodies the decision to neglect spatial correlation.

There are many natural extensions to the modelling we have introduced. We have considered only very simple functional forms (12, 13) for the rate functions, but there is scope (especially given the scale of the data from ray-tracing studies) for exploring much more elaborate forms, or using “non-parametric” methods such as splines. The maximum likelihood framework naturally extends to model selection so criteria such as the likelihood ratio (Cox, 2006), and various information criteria such as Akaike and Bayes (Burnham and Anderson, 2002), each of which is based on the likelihood, can be used for formal comparison between different candidate models. We have focused our attention on direct light, modelling a binary sunlit-shaded state stochastically and the amplitude during sunlit periods by a deterministic light amplitude envelope function. We could similarly model scattered and diffuse light, and thus model the total incident light flux as an additive combination of direct, scattered and diffuse contributions. A limitation of the current work is that we have focused on light dynamics within a static canopy and not yet considered the effects of canopy motion, for example due to wind. Even moderate wind may substantially impact canopy photosynthesis (Burgess et al., 2016) and a goal of ongoing work is to characterise the effect of canopy movement on light dynamics. The technical challenges of imaging and ray-tracing moving canopies are much more substantial (Burgess et al., 2016). However, the mathematical model presented here will, at least as a starting point, be appropriate to apply to ray-tracing data from dynamic canopies once such data are forthcoming. An ultimate goal is to connect the light dynamics to the movement of the canopy, and to connect movement of the canopy to the mechanical properties of the plants comprising it. Achieving this will enable identification of which plant mechanical properties can be targeted for improving biomass production and yield (Burgess et al., 2016).

The heterogeneity of the light environment influences how plants respond to and exploit available resources for photosynthesis and crop production. This has been recently demonstrated using recovery from photoprotection (Kromdijk et al., 2016). However, to quantify the impact of a particular photosynthetic process (Rubisco activation, stomatal opening, photoprotection) on productivity requires knowledge of the precise ‘signature’ of light-shade dynamics. For example, longer periods in high light and low light are likely to be less productive than rapid fluctuations (Roden and Pearcy, 1993; Burgess et al., 2016). This is because rapid fluctuations are likely to result in a higher induction state of photosynthesis. A high induction state means for example that Rubisco is maintained in a high state of activation and stomata remain open for longer. Longer periods of low light cause de-activation of enzymes and stomatal closure. The situation is complex since, for example, stomata that remain open during a period of low light will have a higher transpiration rate and hence their instantaneous water use effciency will be lower during this period (Lawson and Blatt, 2014) despite the enhanced ability to respond to any new high light event. Additionally, the rapid de-activation of non photochemical quenching (NPQ) can be an advantage in low light (Kromdijk et al., 2016) and during the subsequent high light induction (Hubbart et al., 2012). It is only by understanding the precise spatio-temporal light dynamics in different canopy structures, aided by models as we have presented here, that we can predict the impact of these different processes on whole-plant photosynthesis.

## ACKNOWLEDGMENTS

O.E.J., E.H.M., S.P.P. and R.R. were supported by the Biotechnology and Biological Sciences Research Council [BBSRC grant BB/J003999/1]. The authors thank the anonymous reviewers for the suggestions and comments, and Prof Xinguang Zhu and Dr Qinfeng Song (Shanghai Insititue of Plant Physiology and Ecology, Chinese Academy of Sciences) for useful discussions regarding Fast-Tracer.

## APPENDIX

### Direct and diffused light

Solar ray direction depends on the time and location via day, *d*, hour, *t*, and latitude, *l*_*a*_ (Cambell and Normal, 1998). The sun declination angle is

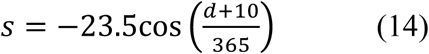

The hour angle is

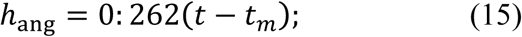

where tm is time of solar noon. The solar elevation angle is

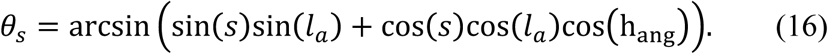

The solar azimuth angle is

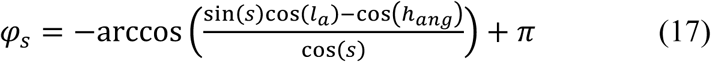

Then the direction of direct light is given by a vector *(x; y; z)* with

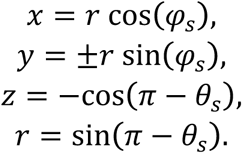

Here *y > 0* if *t<6*hr, and *y* = 0 if *t* = hr.

Direct light irradiance on a unit surface is given by

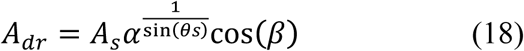

where *α*is atmosphere transmittance, *As*, is a solar constant (2600*µmol m*舑^2^ *s*舑^1^), and *β* is the angle between the light ray and the normal to the surface.

Diffused light irradiance on a horizontal plane is:

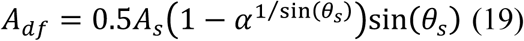

Diffused radiation does not depend on orientations of a leaf (Cambell and Normal, 1998).

### Light response curve

The response of photosynthesis to light irradiance, *A*, is calculated using a non rectangular hyperbola:

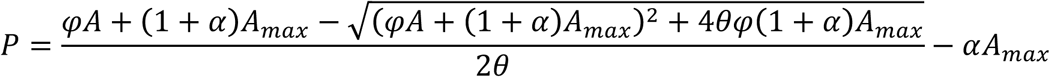

The light response curve is defined by four parameters: the quantum use efficiency *φ*, the convexity *θ*, the maximum photosynthetic capacity, *A*_*max*_, and the rate of respiration, which we assume is proportional to maximum photosynthetic capacity *αA*_*max*_.

